# An Improved Manufacturing Method for Infusible Extracellular Matrix Materials to Enable Efficient Scale-Up, Improved Efficacy, and Clinical Translation

**DOI:** 10.64898/2026.10.29.762959

**Authors:** Michael Nguyen, Emma Zelus, Maria Karkanitsa, Benjamin Liebman, Colin Luo, Carlos Munoz, JuanJuan Liu, Connor Conklin, Pedro Cabrales, Mark Hepokoski, Adam Kinsey, Karen Christman

## Abstract

Decellularized extracellular matrix (ECM) materials are a popular choice for tissue engineering and regenerative medicine applications due to their multifaceted bioactive capabilities arising from their heterogenous composition. For minimally invasive delivery, infusible ECM (iECM) was developed through fractionation of digested decellularized ECM, allowing the material to be safely delivered intravascularly. iECM localizes to permeable vasculature in inflamed tissues and exerts immunomodulatory and pro-reparative effects with proof-of-concept already demonstrated in multiple preclinical models including myocardial infarction (MI) and multiple organ dysfunction syndrome (MODS). However, one major limitation of iECM is the steps required for manufacturing, involving centrifugation, dialysis/lyophilization, and filtration (C-D-F). These steps limit manufacturing output and iECM concentration, and require significant human handling, limiting compliance with current good manufacturing practices (cGMP). To overcome these challenges, we developed a method for manufacturing iECM that utilizes depth filtration and tangential flow filtration (DF-TFF), which reduces manufacturing time, allows for increased iECM concentration, and reduces human manipulation of material, improving translatability and scalability of iECM materials. Based on multiple material characterization methods, we determined that iECM made with DF-TFF was comparable to iECM made with C-D-F. By increasing iECM concentration through this manufacturing method, we demonstrate a dose dependent response in an acute MI porcine model and a mouse MODS model. Finally, DF-TFF enabled timely production of iECM from alternative tissue sources. Altogether, DF-TFF allows for improvements in production time, scale-up, cGMP compliance, and increased dose through higher concentration, paving the way for future clinical translation as well new preclinical studies.

## Introduction

Decellularized extracellular matrix (ECM) materials have seen widespread adoption in the field of tissue engineering and regenerative medicine in recent decades^1^. In particular, ECM materials contain a myriad of bioactive signals from their source tissue, providing pro-reparative and immunomodulatory signals to the area of delivery^2,3^. For these applications, ECM materials have traditionally been used as either implanted materials such as patches^4^, coatings^5^, or hydrogels^3^. While effective, these materials are inherently limited by the need to directly access the tissue of interest, either limiting the material to easy to access tissues, requiring invasive surgery for delivery, or requiring specialized equipment and skill for injection catheter-based delivery^6^.

To overcome these challenges, we recently developed an infusible ECM material (iECM) derived from the sub 220 nm portion of partially pepsin digested myocardium derived ECM^7–11^. These components includes soluble peptides and proteins as well as large macromolecular fibril proteins and polysaccharides originating from their source tissue and generated through partial pepsin digestion^11^. This material is able to be safely delivered into the bloodstream without deleterious coagulation, despite its collagen content^7^, with iECM able to localize to sites of endothelial dysfunction^2,7,9^, specifically in gaps between endothelial cells that arise from inflammation^7^. We previously tested iECM in small and large animal models of myocardial infarction (MI)^7,10,11^, as well as small animal models of traumatic brain injury (TBI)^8^ and multiple organ dysfunction syndrome (MODS)^9^. Once delivered, iECM reduces vascular permeability and exerts immunomodulatory effects on surrounding tissue, resulting in improvements in cardiac function in the case of MI^7^ and survival in the case of MODS^9^. Altogether, iECM has the potential for use in many conditions characterized by inflammation and increased vascular permeability while being able to be delivered minimally invasively.

To fabricate iECM, we previously employed a method based on centrifugation, dialysis/lyophilization, and filtration (C-D-F)^7^. After digestion, centrifugation was employed to remove most of the undigested larger particulate matter based on density, with the lighter supernatant being concentrated through desalting with dialysis, lyophilization to remove water, and weight-based resuspension in appropriate salt concentrations. Following reconstitution, the solution was then membrane filtered to remove large components that the centrifugation could not remove before being filtered through a 220 nm filter for sterilization. While this C-D-F method yielded sufficient quantities of material for material characterization and initial preclinical animal testing, the C-D-F manufacturing method contained several inherent limitations that impeded clinical translation of iECM. First, the concentration through dialysis and lyophilization required multiple days to complete. Additionally, iECM final concentration was limited by the final membrane filtration used for further separation of large components, as too high of a pre-filter solution concentration would be unable to pass through the filter. This in turn limited testing of iECM at higher concentrations to observe potential enhancements in therapeutic efficacy. Finally, the filling and unloading dialysis tubing also required significant open-air human handling of the solution, introducing multiple points of potential contamination. This limits transfer to a current good manufacturing practice (cGMP) environment, limiting its clinical translation^12,13^.

To overcome these limitations, we developed a new iECM manufacturing method that adapted manufacturing practices currently used for large scale production of other biologics^14^ and applied them to the separation and concentration of infusible ECM components. First, we utilized body feed depth filtration for the clarification of the digested ECM solution through the removal of large ECM particulate matter. Following filtration, we employed tangential flow filtration (TFF) to concentrate iECM to therapeutically relevant concentrations. This novel ECM processing method employing depth filtration and TFF (DF-TFF) significantly reduces preparation time compared to our original method, allows for production of higher concentration iECM through improved separation efficiency, and allows for processing in a closed system after filtration, reducing the potential for contamination. In this work, we validate our improved method for iECM production by assessing the material characteristics and efficacy through biochemical analysis, *in vitro,* and *in vivo* testing. We further demonstrate the utility of the DF-TFF iECM production method by expanding iECM manufacturing to alternative tissue sources.

## Results

### 2.1: Development of New iECM Manufacturing Processes

To produce decellularized ECM for iECM manufacturing, porcine left ventricular myocardium was first minced and decellularized with 1% sodium dodecyl sulfate (SDS). After decellularization, tissue was then lyophilized, milled into a fine powder, and digested with pepsin (Fig S1a, S2a). The isolation and preparation of the infusible portion (components smaller than 220 nm) of digested decellularized ECM involves two major requirements: the removal of large particulate matter and the concentration of isolated ECM products to therapeutically relevant concentrations. In our original C-D-F method (Fig 1a), high speed centrifugation (RCF = 15000 g) was first used to pellet out most of the large particulate matter based on density, leaving behind a supernatant that was mostly comprised of the smaller digested ECM products that would become iECM (Fig S1b). However, in order to achieve sufficient pelleting, centrifugation needed to be performed for extended period of time, taking on average 3 hours to process 1g of digested ECM. These components were then concentrated through dialysis (Fig S1c) and lyophilization (Fig S1d), where the recovered ECM components were desalted, dried, and then reconstituted at a higher concentration in PBS (Fig S1e). To reach sufficient removal of salts, dialysis needed to be performed for at least 48 hours, with the lyophilization of the desalted ECM taking on average 120 hours with a lab scale lyophilizer. Centrifugation was unable to remove all large particulate matter, so a secondary separation membrane filtration was necessary prior to terminal sterile filtration (Fig S1f). Because of the buildup of residual large ECM particulate, separation filtration took on average 4 hours to complete. Altogether, starting from the decellularized ECM powder, this process took approximately 223 hours in total, with the bulk of the time due to dialysis and lyophilization for iECM concentration.

**Figure 1:**
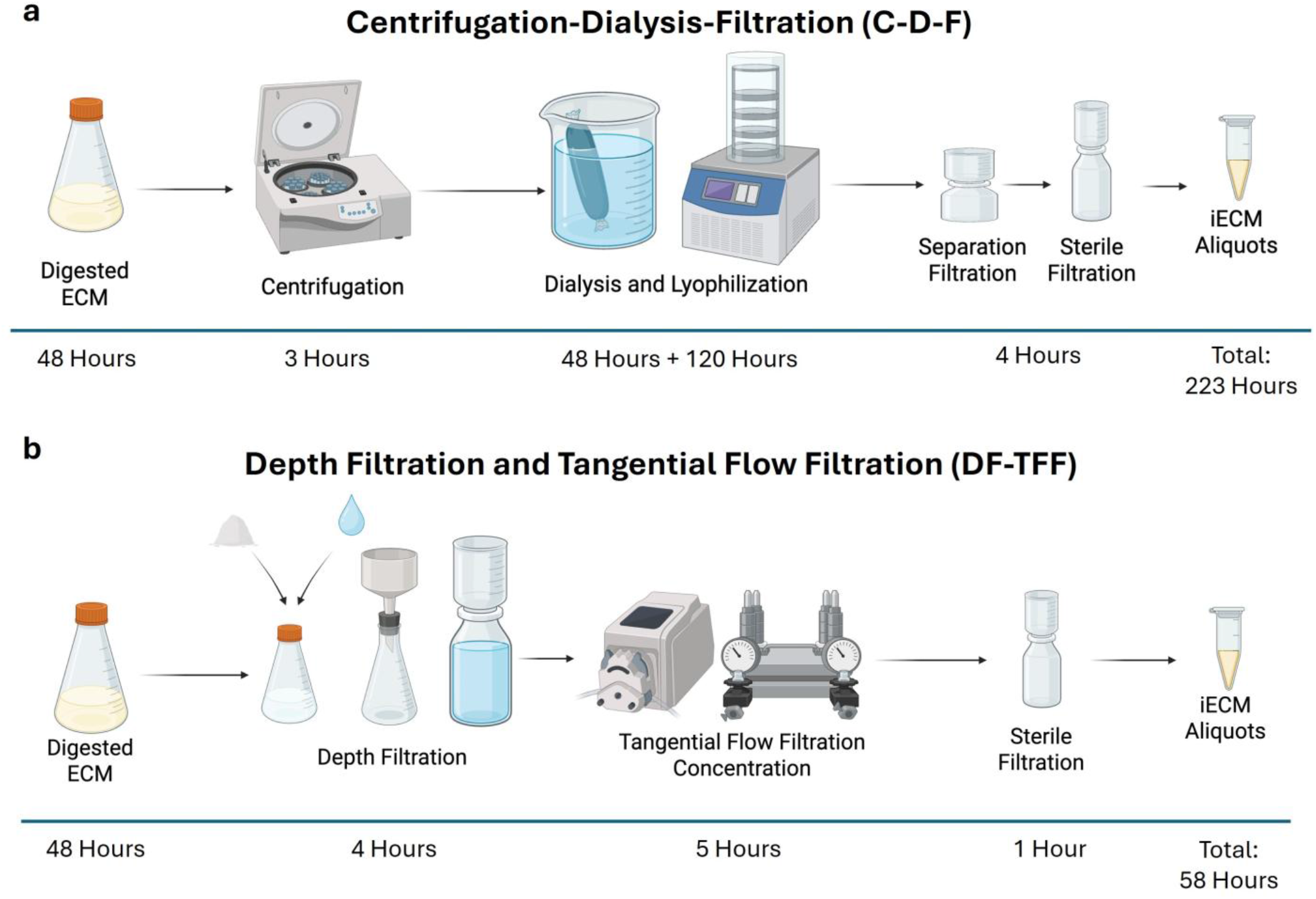
Processes used to manufacture iECM from digested, decellularized ECM. a) Original iECM manufacturing method involving centrifugation, dialysis, and filtration (C-D-F) with approximate time required to process 1 g of decellularized ECM. b) Optimized iECM manufacturing method using depth filtration and tangential flow filtration (DF-TFF) with approximate time required, demonstrating significant reduction in time required to process 1 g of decellularized ECM powder.

In our new DF-TFF method (Fig 1b), removal of large particulate matter from decellularized and digested ECM is now achieved through body feed depth filtration, using pharmaceutical grade diatomaceous earth as the filter aid. This creates a three-dimensional filter that is able to more efficiently remove the range of undesired large particulate material prior to concentration^15^. The digested ECM was diluted and combined with diatomaceous earth. The resulting slurry was then filtered through a 20 μm filter (Fig S2b), where the diatomaceous earth particle size keeps it retained during filtration. As the filter aid settles, the particles form a deep porous filter cake that traps large particulates^15^, including the undigested ECM particulate, while allowing smaller components to pass around, maintaining a timely filtration rate. This is in contrast with membrane filtration where large particulates would build up at the membrane surface and reduce filtration speed. While a secondary membrane filtration was performed after depth filtration, this was primarily done to ensure all of the filter aid was removed and did not significantly contribute to processing time. Through optimization of iECM filtration concentration and filter aid feed ratio, we were able to complete the removal of large ECM particulate in 4 hours. For material concentration, TFF against a 1 kDa molecular weight cutoff membrane wawas used to concentrate ECM digestion products (Fig S2c). TFF has previously been used in the production of biologics and extracellular vesicles, where maintaining buffer conditions is critical for preserving the qualities of the product^16^. As such, we leveraged this technology to preserve the salt concentration while concentrating iECM, removing the need for desalting as was used in the C-D-F method. This method of iECM concentration took on average 5 hours, a significant reduction in time compared to the dialysis and lyophilization previously used. With this new method, 1 g of decellularized ECM could now be processed into iECM in 58 hours, with the separation and concentration now only taking an additional 10 hours on top of the standard 48 hour pepsin digestion. Though different in their approach and associated required time, both of the C-D-F and DF-TFF methods process digested decellularized ECM to where it can be filtered through a 220 nm filter, resulting in a sterile ECM product that can be safely intravascularly infused.

### 2.2: Comparison of iECM material properties between manufacturing methods

Given we had already performed extensive studies showing efficacy of iECM^7–9,11,17^, including a good laboratory practice (GLP)-like porcine MI study, we wanted to confirm that the iECM made using the DF-TFF remained equivalent to iECM made with the previous C-D-F method in anticipation of an investigational new drug (IND) submission, we first explored whether key material properties were altered through the change in processing method. These included the protein/peptide size distribution, safety criteria related to material injectability, and biochemical properties. Because ECM materials are inherently heterogenous, multiple batch replicates were made using the C-D-F and DF-TFF methods to account for any batch variability.

To investigate potential effects on protein/peptide size distribution, we tested multiple batches of both C-D-F and DF-TFF iECM using SEC. When analyzed using size exclusion chromatography (SEC) (Fig 2a,b), the peaks of C-D-F and DF-TFF iECM are consistent with each other: a high molecular weight (MW) peak (apparent MW = 1900 kDa), a medium MW peak (apparent MW = 77 kDa), and a low MW peak (apparent MW = 3 kDa) (Fig 2a). When each peak was analyzed for their amount relative to the total protein/peptide content, each MW peak was found to be equivalent between C-D-F and DF-TFF iECM (Fig 2b), suggesting the DF-TFF manufacturing method did not significantly alter the protein/peptide distribution of iECM.

**Figure 2:**
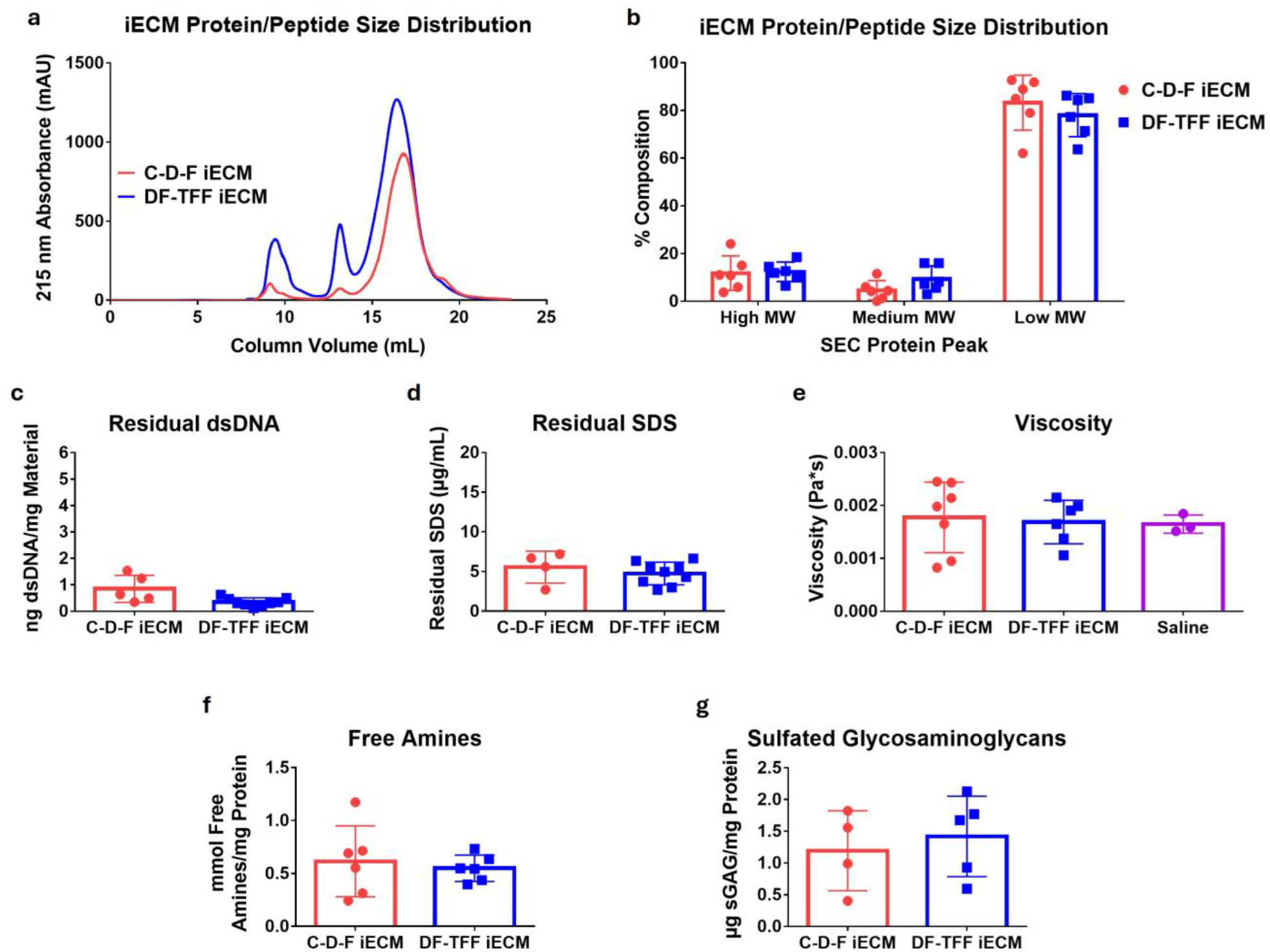
Characterization and comparison of material properties of iECM made with C-D-F and DF-TFF methods. a) Average SEC trace showing comparable protein/peptide distribution of iECM batches between the two manufacturing methods (n=5 C-D-F and DF-TFF). b) Quantification of relative distribution of iECM batches separated into high MW, medium MW, and low MW peaks (n=5 C-D-F and DF-TFF). c-e) Safety metrics related to infusibility showing comparable c) residual dsDNA (n=5 C-D-F, n=9 DF-TFF), d) residual SDS (n=4 C-D-F, n=9 DF-TFF iECM), and e) viscosity between iECM from both manufacturing methods (n=7 C-D-F, n=6 DF-TFF, n=3 saline). f-g) Biochemical analyses showing comparable f) free amine (n=6 C-D-F and DF-TFF) and g) sGAG content (n=4 C-D-F, n=5 DF-TFF) between iECM manufacturing methods. All metrics showed no statistically significant difference between groups, demonstrating equivalence between C-D-F and DF-TFF iECM. Statistical significance (p < 0.05) determined by unpaired t-test for comparisons of two groups and a one-way ANOVA for comparisons between three groups.

Next, to ensure that DF-TFF iECM remained safe for intravascular infusion, we assessed several quality control parameters associated with iECM: residual dsDNA^18^, residual sodium dodecyl sulfate (SDS)^19^, and viscosity^20^. The presence of dsDNA in iECM can originate from incomplete decellularization or contamination during the manufacturing process, where the C-D-F method is more susceptible to contamination due to the human manipulation of liquid during dialysis bag filling. However, both C-D-F and DF-TFF iECM had low dsDNA values, suggesting that neither manufacturing process was introducing contamination (Fig 2c). For SDS, which is used to decellularize the starting tissue, iECM has been found to have lower residual SDS compared to the full ECM digest it is derived from^7^. It was previously thought that the low residual SDS content was in part due to the complete desalting step of the C-D-F manufacturing method, where residual SDS would diffuse out of the dialysis tubing. However, low residual SDS was found in both C-D-F and DF-TFF iECM, suggesting that the reduction in residual SDS from the ECM digest was inherent to the ECM separation, both when centrifugation and depth filtration are used (Fig 2d). Finally, the average viscosity of iECM batches made with the C-D-F and DF-TFF methods was found to be comparable to saline, confirming that it was suitable for intravascular infusion (Fig 2e). Altogether, these data suggest that the DF-TFF manufacturing process did not significantly alter infusibility of iECM compared to previously produced batches.

Finally, we compared the free amines and sulfated glycosaminoglycans (sGAGs) present in iECM made with both manufacturing methods. In this context, free amines represent a complementary measure of protein/peptide content and serve as readily available moieties for chemical modification of iECM, particularly for fluorescent labeling of iECM^7–9^. Furthermore, free amines serve as a rough measure of ECM fragmentation by pepsin, as enzymatic cleavage of peptide bonds contributes to a higher free amine content. sGAGs can impart bioactivity through their ability to bind many growth factors^21,22^. While the myocardial ECM hydrogel form was shown to have more sGAGs, iECM did contain small amounts^7^. When normalized to the average protein content, iECM from both methods had equivalent amounts of free amines (Fig 2f). Similar trends were observed with sGAG content of iECM batches, where similar amounts of sGAGs were detected between iECM of both manufacturing methods (Fig 2g). These data, compounded with protein/peptide distribution data, suggest that the composition of iECM was not affected by the DF-TFF manufacturing method.

### 2.3: Effect of iECM manufacturing method on cytocompatibility and hemocompatibility

Following biochemical characterization, we wanted to ensure that DF-TFF iECM remained cyto- and hemocompatible. We first tested cytocompatibility of DF-TFF iECM and C-D-F iECM by incubating primary rat coronary artery endothelial cells, the cell type that would readily interact with iECM following intravascular delivery and binding to permeable vasculature^7^. No batch of iECM had negative effects on cell survival, demonstrating that the improved manufacturing method did not introduce components that would negatively impact cells (Fig 3a).

**Figure 3:**
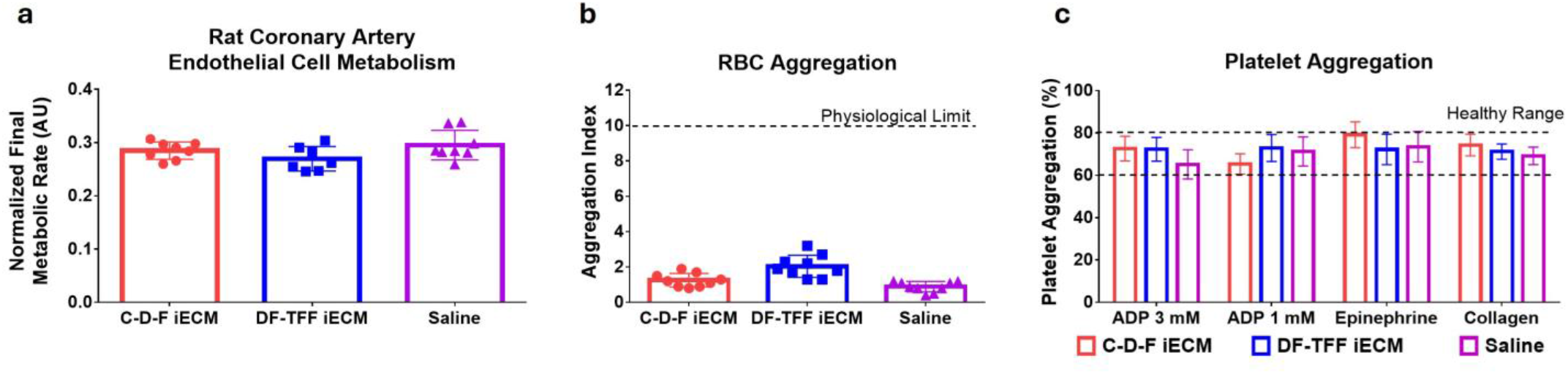
Cytocompatibility and hemocompatibility of iECM batches. a) Cytocompatibility tests with rat coronary artery endothelial cells showing equivalent cytocompatibility of iECM from both manufacturing methods compared to saline treated cells (n=8 C-D-F, n=7 DF-TFF, n=8 saline). b) RBC aggregation when whole blood is mixed with C-D-F iECM, DF-TFF iECM, or saline (n=9 C-D-F, DF-TFF, and saline), indices below physiological limits. c) Effect of iECM batches or saline on platelet aggregation (n=4 C-D-F, DF-TFF, and saline per agonist), showing platelet aggregation remains in healthy ranges when treated. Statistical significance (p<0.05) determined with one-way ANOVA.

We then tested the hemocompatibility of DF-TFF iECM compared to C-D-F iECM, which is critical given the intravascular delivery modality. With regard to red blood cell (RBC) aggregation, DF-TFF iECM did not significantly increase RBC aggregation over C-D-F iECM or a saline control, with all groups falling within physiologically safe RBC aggregation indices (Fig 3b)^23,24^. Similar results were found with platelet aggregation, where platelets stimulated with multiple activators did not show differences in aggregation activity when mixed with iECM batches or saline (Fig 3c)^24^. These results, combined with viscosity data, demonstrated that iECM manufactured with the DF-TFF method remained safe for intravascular delivery.

### 2.4: Effect of iECM manufacturing method on retention in a rat model of MI

Previous work suggested that iECM likely localizes to sites of endothelial dysfunction in inflamed tissues through specific binding to exposed basement membrane ECM proteins or cell surface receptors^11^. Because this localization may be sequence specific, we wanted to assess whether this binding capacity was retained when iECM was produced using the DF-TFF method. This was assessed in a rat model of MI as previously described^7,11^. One hour following administration, which was immediately after reperfusion through a simulated intracoronary infusion, hearts were harvested, sliced, and scanned with near infrared imaging. The fluorescent signal in the infarcted regions of the heart was found to be the same for both C-D-F and DF-TFF iECM (Fig 4a,b). This suggests that the production of iECM with the DF-TFF method did not significantly affect the binding capability of iECM in inflamed tissues.

**Figure 4:**
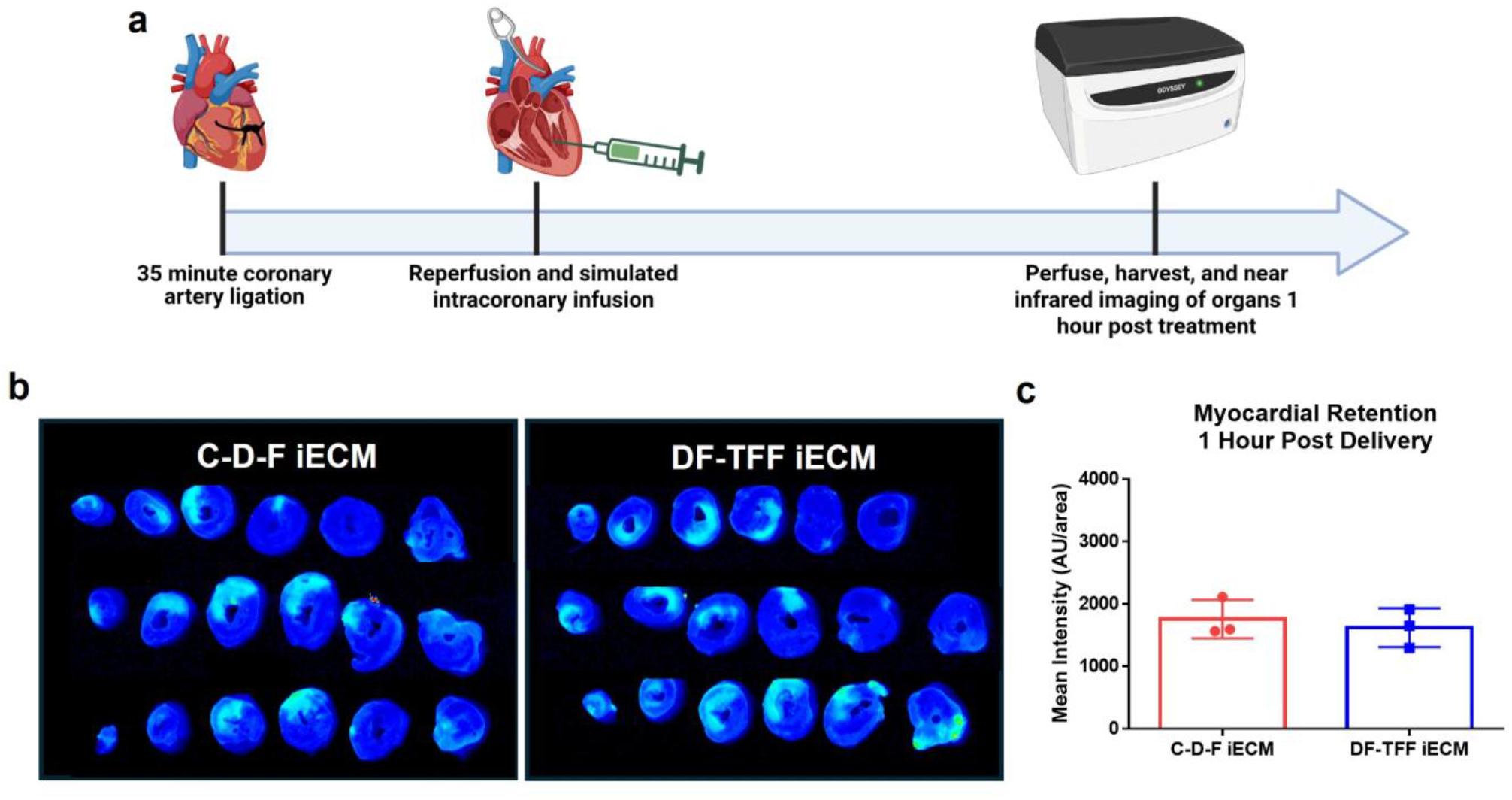
Retention of iECM in a rat MI model. . a) Timeline of iECM retention experiment in rat MI model. b) Near infrared images of transverse heart sections from rats treated with C-D-F iECM or DF-TFF iECM harvested 1 hour post reperfusion showing infarct targeting. b) Quantification of infarct near infrared fluorescent signal demonstrating equivalent retention of C-D-F and DF-TFF iECM following MI and reperfusion (n=3 C-D-F and DF-TFF). Statistical significance (p < 0.05) was determined with an unpaired t-test.

### 2.5: The DF-TFF iECM manufacturing method enables increased production scale and increased iECM concentration

Due to the significant improvements in processing time and the scalable nature of DF and TFF, we next assessed the effects of increased manufacturing scale on iECM material properties, comparing iECM from 1g and 4g digests processed using the DF-TFF method. Because of the viscosity of the digested ECM solution, it was expected that an increase of the mass of ECM and volume of digestion would alter the digestion conditions given the effect solution viscosity has on enzyme kinetics^25,26^. This in turn could potentially result in alterations to the protein/peptide distribution of iECM batches. Similarly, processing increased volumes of digested ECM through the DF-TFF method may cause unexpected changes in composition. As such, we needed to ensure that the switch from processing 1g of decellularized ECM to 4g did not significantly alter the properties of iECM. The primary readout for this assessment was the protein/peptide distribution as determined by SEC. When averaged across multiple batches of iECM produced from 1g digests and 4g digests, the same major peaks were detected (Fig 5a). Furthermore, the proportion of these peaks relative to total protein/peptide content was consistent between production methods (Fig 5b), suggesting that the DF-TFF method could allow for scaled-up manufacturing of iECM without significant change to the protein/peptide composition.

**Figure 5:**
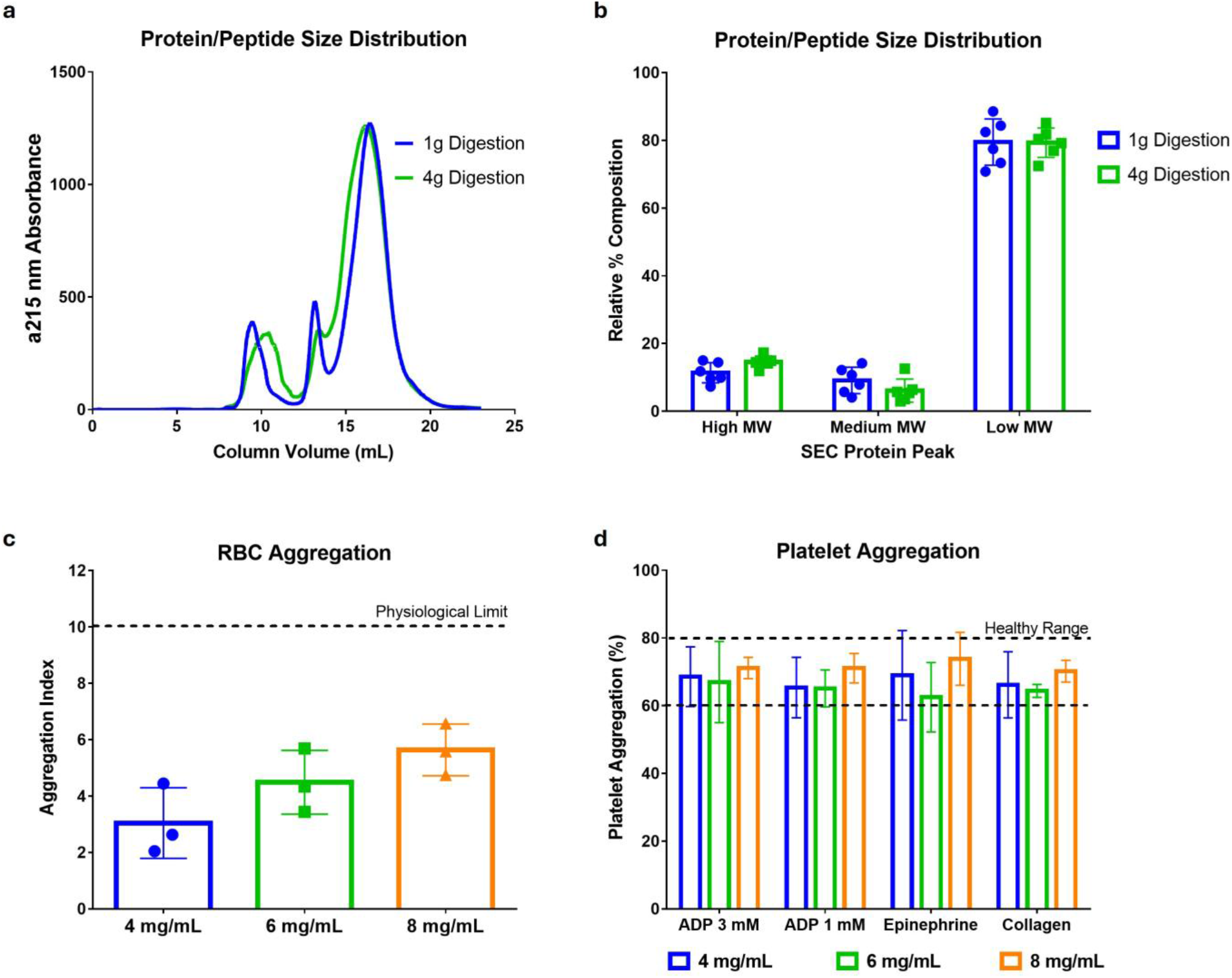
Characteristics of iECM batches based on digestion scale. a) Average SEC trace of iECM batches made with 1g digests (n=6) and 4g digests (n=6) showing comparable peak patterns between digest scales. b) Quantification of relative area of major peaks for 1g (n=6) and 4g (n=6) digestion iECM batches showing comparable distribution of protein/peptide sizes. c) Effects of iECM concentration on RBC aggregation (n=3 for 4 mg/mL, 6 mg/mL, and 8 mg/mL), with all concentrations resulting in aggregation indices below physiological limits. d) Effects of iECM concentration on platelet aggregation (n=3 for 4 mg/mL, 6 mg/mL, and 8 mg/mL per agonist), showing platelet aggregation within healthy limits. Statistical significance (p<0.05) determined with multiple t tests for iECM SEC peaks, one-way ANOVA for RBC aggregation, and multiple one-way ANOVAs for each platelet aggregation agonist.

Previously, the C-D-F method limited iECM concentration to 4 mg/mL due to the separation efficiency of centrifugation. Because a secondary filtration was required post concentration, iECM had to be resuspended at a higher concentration to account for large particulate removal, and because membrane filtration was used, increased concentrations of iECM were difficult to filter due to clogging of the filter. In contrast, depth filtration of the digested ECM resulted in significantly improved separation of large ECM particulate, allowing control over iECM concentration through TFF without the need for a secondary separation filtration prior to terminal sterile filtration. To ensure that the higher concentration of iECM did not affect its hemocompatibility, we assessed the effects of 4, 6, and 8 mg/mL iECM on RBC and platelet aggregation. While trending increases in RBC aggregation were observed (Fig 5c), these values stayed within physiological ranges^23,24^. Similarly, all platelet aggregation values were within healthy ranges^23^ when blood was treated with iECM of increasing doses (Fig 5d). These data suggest that increased iECM concentration could be safely delivered without hemocompatibility issues, allowing us to further probe the effects of increased iECM dosing.

### 2.6: Assessment of DF-TFF iECM in a porcine acute MI model

Leveraging improvements in manufacturing output and the potential for increased iECM concentration afforded by the DF-TFF process, we assessed the effect of increasing iECM doses in a porcine acute MI model. While our previous investigations looking at iECM in acute MI preclinical models were limited to 4 mg/mL doses^7^ due to inherent C-D-F process attributes, we were now able to assess whether increased iECM dosing led to increases in iECM retention and early improvements in cardiac function with increasing iECM dose. To assess these potential changes, we assessed iECM in a porcine acute MI model, treating pigs with 4 mL of saline or 4 mL of fluorescently labeled 4, ,6, or 8 mg/mL iECM through an intracoronary infusion immediately after reperfusion (Fig 6a). Pigs were euthanized 24 hours after reperfusion to evaluate both retention and early changes in cardiac function. Prior to intracoronary infusion, echocardiography was performed post-MI to ensure adequate infarcts and then again at 24 hours after reperfusion to assess changes in myocardial wall motion and cardiac function. After euthanasia, hearts were sliced into transverse sections and scanned using near infrared imaging to assess the presence of iECM in the myocardium. In this case, no significant difference in signal was found between iECM of any dose, with all having significantly higher signal in the infarcted tissue compared to saline treated animals (Fig S3).

**Figure 6:**
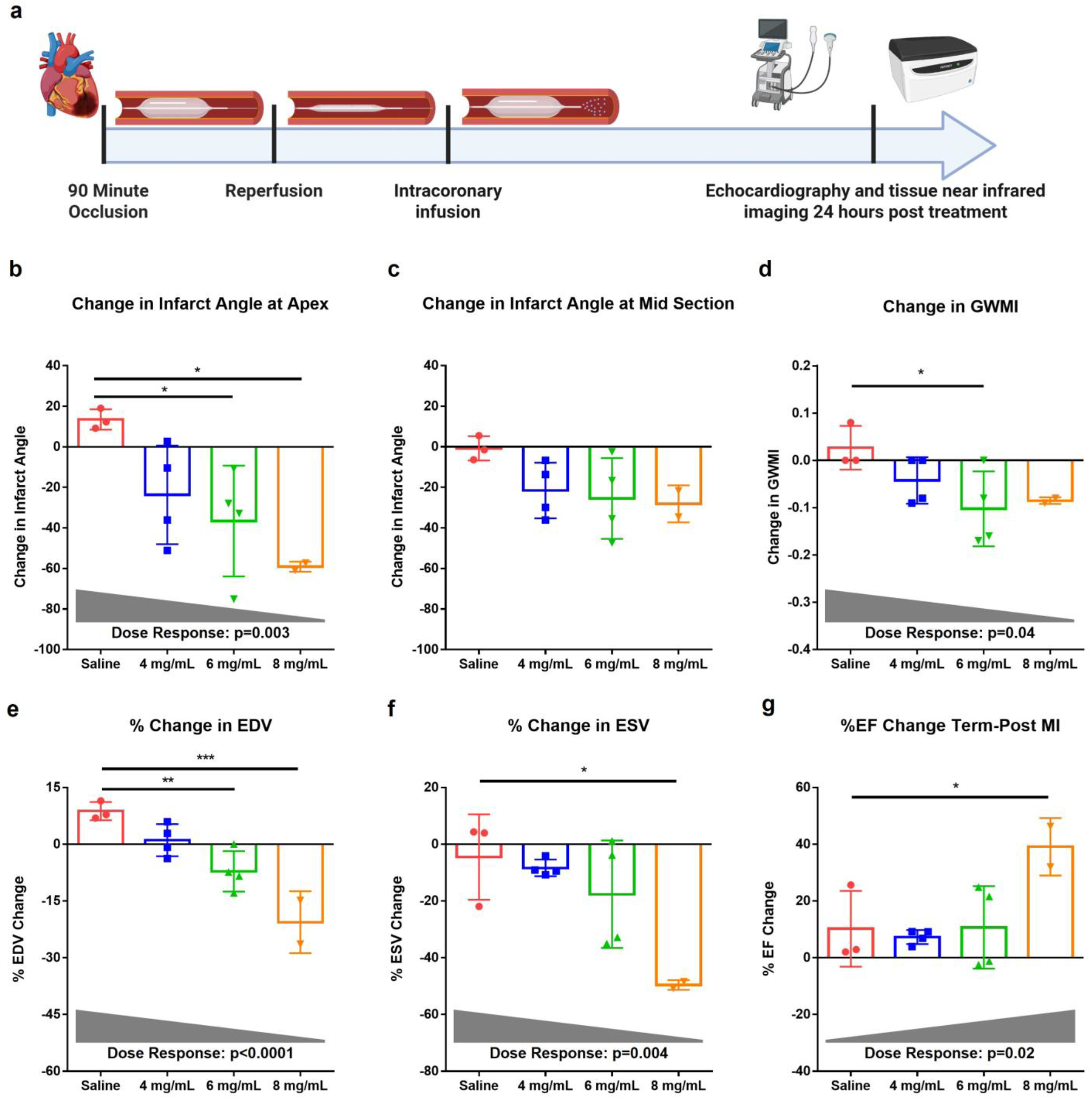
Dose escalation effects of DF-TFF iECM on retention and cardiac function in a porcine acute MI model. a) Timeline of experiments for testing effects of iECM dose concentration in a porcine acute MI model. Following anesthesia, pigs underwent MI through the inflation of a balloon catheter in the left anterior descending artery. After 90 minutes of ischemia, balloons were deflated to allow for reperfusion, with a post-MI echocardiographic assessment of myocardial wall motion and cardiac function performed. Balloons were then reinflated for intravascular administration of saline or iECM under stop-flow conditions, then deflated again post infusion. 24 hours post treatment, a terminal assessment of myocardial wall motion and cardiac function was performed. b-d) Reductions in wall motion abnormalities determined by echocardiography to quantify b) changes in the infarct angle at the apex, c) changes in the infarct angle at the mid section, and d) changes in GWMI at 24 hours post-treatment compared to immediately after reperfusion (post-MI and pre-treatment). Changes in apex infarct angle and GWMI demonstrated significant (p<0.05) dose dependent responses. e-g) Improvements in cardiac function as determined by echocardiography to quantify % changes in e) EDV, f) ESV, and g) EF at 24 hours post treatment compared to immediately after reperfusion. All cardiac function metrics demonstrated significant (p<0.05) dose dependent responses. Statistical significance determined with one-way ANOVA with post-hoc Dunnett test compared to saline (*p<0.05, *p<0.01, **p<0.001). Significant (p<0.05) dose response of iECM was determined with a post-hoc linear trend analysis. (n=3 saline, n=4 4 mg/mL and 6 mg/mL, n=2 8 mg/mL)

To ensure that the increases in iECM concentration did not induce adverse effects following intravascular administration, blood was also collected at euthanasia for evaluation of hematology, serum chemistry, and coagulation parameters (Fig S4). Blood urea nitrogen (Fig S4a) and creatinine levels (Fig S4b) in pigs treated with iECM were nonsignificant compared to saline, indicating kidney function was not impaired. Similarly, blood bilirubin levels (Fig S4c) in iECM treated animals remained consistent with saline treated animals, indicating no liver toxicity. Furthermore, blood cell counts, including total white blood cell (Fig S4d), red blood cell (Fig S4e), platelets (Fig S4f) and neutrophils (Fig S4g) remained consistent between saline and iECM treated animals. Finally, blood clotting metrics including activated partial thromboplastin time (Fig S4h) and prothrombin time (Fig S4i) were unaffected by iECM treatment, indicating increased iECM content also did not impact blood clotting. Altogether, these data indicate increased iECM concentration did not induce adverse effects.

Following ischemia and reperfusion, the injured myocardium contracts abnormally due to cardiomyocyte death and tissue dysfunction^27^. To probe the effects of iECM concentration on cardiac function at early timepoints, we observed changes in wall motion abnormality at 24 hours post-treatment relative to immediately after reperfusion (post-MI and pre-treatment). First, by measuring the angle of abnormal left ventricular free wall motion, we were able to estimate the area of the infarcted tissue^28^, with this measurement being performed at the apex of the heart and at the midsection. When treated with saline, infarct angle increased over time at the apex (Fig 6a) and remained constant at the midsection of the heart (Fig 6b). When treated with iECM, a significant linear dose response (p = 0.003) was seen with reductions in apex infarct angle as iECM concentration increased (Fig 6a). The mean infarct angle at the midsection also decreased with iECM treatment on average with all treatments having the same degree of change, but this was not statistically significant (Fig 6b). In this model, infarcts caused by the balloon angioplasty mediated occlusion tended to be more apical, resulting in more prominent improvements in apex infarct angle compared to at the mid-section. When looking at the global wall motion index (GWMI), the aggregate index of all wall motion scores^29^, increased iECM concentration also demonstrated a significant dose dependent linear trend in GWMI improvements (p = 0.04), with the 6 mg/mL dose resulting in a significantly larger reduction in GWMI compared to saline (Fig 6c).

While changes in left ventricular (LV) volume and ejection fraction usually manifest after significant negative LV remodeling, early changes in these metrics can indicate preservation of long-term cardiac function^30^. When looking at cardiac function metrics through echocardiography, reductions in end diastolic volume (EDV) compared to post-MI linearly followed increased iECM concentrations (p<0.0001), with 6 and 8 mg/mL iECM having significant reductions compared to saline (Fig 6e). Similarly, increases in iECM concentration also followed a significant linear dose response (p = 0.004) regarding reductions in end systolic volume (ESV), with pigs treated with 8 mg/mL iECM showing significant reductions in ESV compared to saline. Finally, improvements in ejection fraction (EF) also followed a significant linear trend with increasing iECM concentration (p = 0.02), with pigs treated with 8 mg/mL iECM also showing significant improvements in EF compared to saline treated animals. Altogether, these data demonstrate that increasing iECM concentration, which was enabled by the new DF-TFF manufacturing method, led to a dose dependent improvement in cardiac wall motion abnormalities and cardiac function.

### 2.7: Effect of iECM manufacturing method on bioactivity in a mouse MODS model

In multiple models of inflammation, iECM has been shown to reduce inflammation induced vascular permeability and exert immunomodulatory effects, with this phenomenon observed in rodent models of MI^7,11^, TBI^8^, and MODS^9^. In these conditions, acute inflammation causes disruptions in endothelial tight junctions, resulting in dysregulation of fluid flow across the endothelium and into the tissue^31^. This endothelial dysfunction can allow for increased immune cell extravasation and cytokine diffusion, exacerbating inflammation in the tissue^32^. Based on the results from the iECM dose escalation study in pigs, we wanted to compare an increased dose (6 mg/mL) of iECM made with the DF-TFF process to the original 4 mg/mL C-D-F iECM, which was previously evaluated in a mouse MODS model (Fig 7a)^9^. Using these two treatments, we studied changes in vascular permeability and the expression of *Il6* in the lungs, with *Il6* serving as a marker of inflammation^33^.

**Figure 7:**
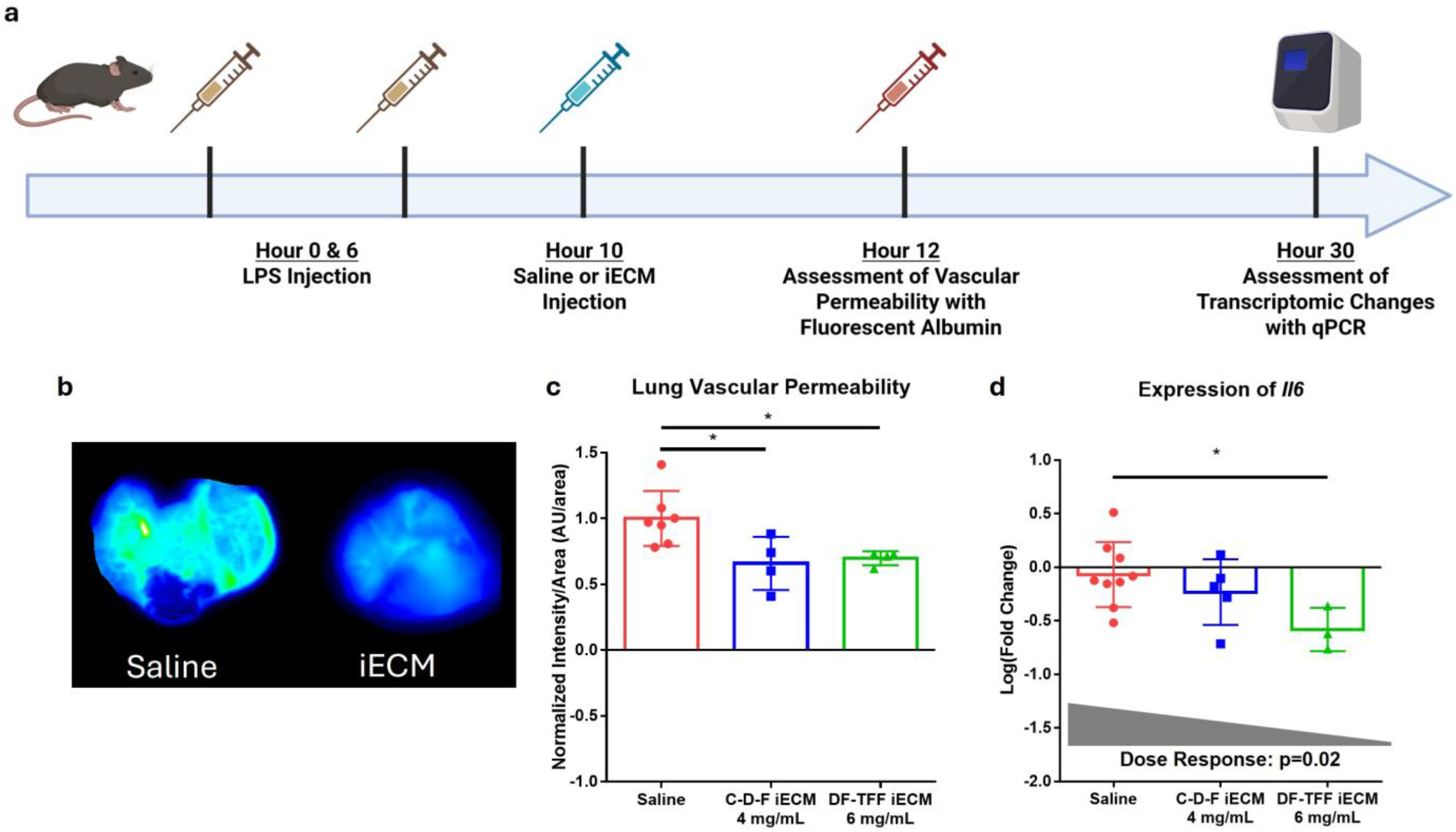
Effect of iECM dose on reductions in vascular permeability and *Il6* expression in a mouse MODS model b. a) Timeline of mouse MODS model showing dual LPS ose to induce a hyperactive immune response characteristic of MODS. 10 hours after initial LPS dose, iECM or saline is administered. 12 hours later, vascular permeability is determined with the administration of a fluorescent BSA tracer, 30 hours later, mice lungs are processed for assessment of *Il6* expression. b) Representative images of mouse lungs after saline (left) and DF-TFF iECM (right) treatment. c) Reductions in vascular permeability in mouse lungs when treated with iECM as determined by reductions in fluorescent BSA signal. (n=7 saline, n = 4 C-D-F 4 mg/mL and DF-TFF 6 mg/mL) d) Downregulation of *Il6* expression in mouse lungs when treated with increasing concentrations of iECM (n=9 saline, n=5 C-D-F, n=3 DF-TFF). Statistical significance (p<0.05) determined with one-way ANOVA with post-hoc Dunnet test compared to saline (*p<0.05).

To simulate the systemic hyperactive immune response that results in MODS, we performed repeated intraperitoneal injections of lipopolysaccharide (LPS), with iECM administered through tail vein injection 10 hours after the initial LPS dose (Fig 7a) as we previously performed^9^. Vascular permeability was assessed through a second tail vein injection of a fluorescently labeled bovine serum albumin (BSA) tracer 1 hour after iECM administration, with organs harvested and the presence of BSA determined using near infrared scanning. For *Il6* expression, mice underwent the same LPS and iECM administration regime, with lungs harvested 30 hours after the initial LPS dose and processed to extract RNA for qPCR.

When looking at vascular permeability, intravascular delivery of either 4 mg/mL C-D-F and 6 mg/mL DF-TFF iECM resulted in significant reductions in lung vascular permeability compared to the saline control as determined by the reductions in extravasation of fluorescently labeled BSA tracers (Fig 7a,b). When assessing *Il6* gene expression in the lungs, the degree to which *Il6* was downregulated followed a linear dose response (p = 0.02), with 6 mg/mL DF-TFF iECM significantly reducing *Il6* expression compared to saline treated animals. These data indicate that increases in iECM dose resulted in improved immunomodulation, corroborating the dose response data obtained from the porcine model of MI.

### 2.8: Application of DF-TFF iECM manufacturing method to alternative decellularized tissues

To further demonstrate the efficiency and reproducibility of iECM manufacturing through the DF-TFF method, we manufactured iECM from alternative tissues: decellularized porcine lung, aorta, and skeletal muscle (Fig 8). Previous data has shown that ECM biomaterial bioactivity depends on the source tissue, as the different protein, peptide, and polysaccharide composition can result in different biological effects^5,34^. With the C-D-F method, only one batch could be made at a given time due to the required space for dialysis and lyophilization. As such, determining properties specific to iECM’s source tissue would have been difficult due to the long production times required for a single batch, with multiple batches required to differentiate between batch specific and tissue specific differences. However, with the improved production speed and scalability afforded by the DF-TFF method, iECM from alternative tissue sources could be made in a fraction of the time previously required.

**Figure 8:**
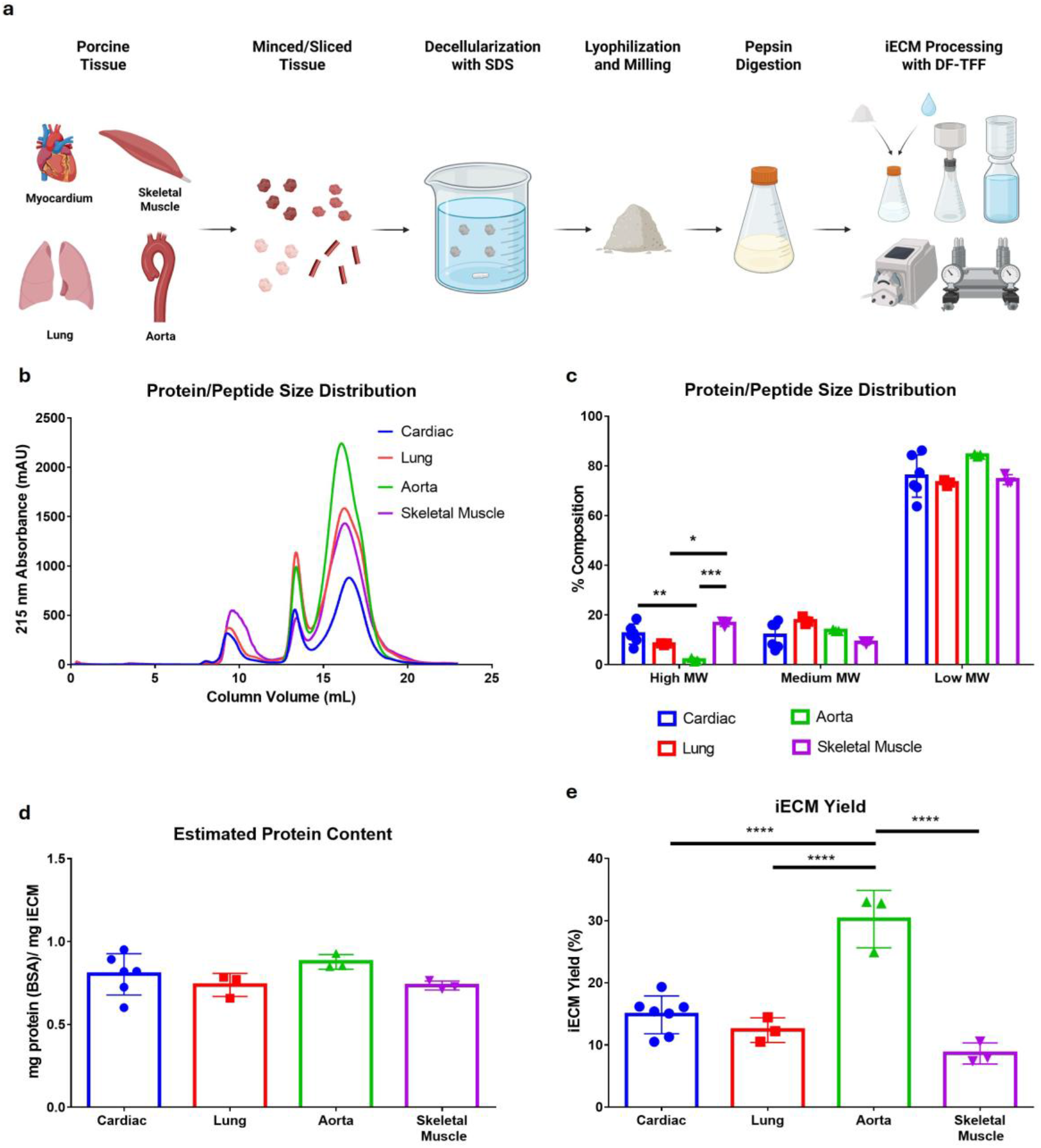
Protein characteristics of iECM manufactured from alternative decellularized tissue sources. a) Processing of alternative tissues for iECM production. Porcine myocardium, lung, skeletal muscle, and aorta tissue were minced or sliced, and then decellularized with SDS. Decellularized tissues were then lyophilized and milled for a 48 hour pepsin digestion. Digested tissues were then processed into iECM using the DF-TFF method. b) Average SEC trace of cardiac, lung, aorta, and skeletal muscle derived iECM batches showing comparable peak locations. b) Quantification of relative peak area of major peaks showing differences in high MW content. cc) Estimated protein content of iECM based on tissue source based on 215 nm absorbance. dd) Average yield as a percent mass recovered from initial decellularized ECM powder of iECM batches showing differences based on source tissue. Statistical significance was determined by one-way ANOVA with post-hoc Tukey test (*p<0.05, **p<0.01, ***p<0.001, ****p<0.0001) (n=6 cardiac iECM, n=3 lung, aorta, and skeletal muscle iECM

Similar to cardiac iECM, porcine lungs, skeletal muscle, and aorta were prepared by mincing, decellularizing with SDS, lyophilizing, and milling prior to a 48 hour pepsin digestion (Fig 8a). While cardiac, lung, and skeletal muscle could be minced using a meat grinder, the high elasticity of aortic tissue necessitated slicing by hand. However, following decellularization and lyophilization, all tissues behaved similarly with regard to milling and digestion. 1 g of digested ECM was then processed through DF-TFF, with all digested ECM solutions being amenable to the DF-TFF process. All batches of iECM from the different tissue sources followed the trimodal behavior seen with cardiac derived iECM, suggesting this protein/peptide distribution pattern is consistent between SDS decellularized ECM that undergoes pepsin digestion (Fig 8b). However, the relative ratio of these three components differed slightly based on tissue source (Fig 8c). This was primarily seen in the differences in high MW content, where aorta derived iECM had significantly less high MW compared to all other batches (Fig 8c). While all batches had similar estimated protein/peptide contents as determined by a BSA standard, regardless of tissue source (Fig 8d), the reduced high MW content appeared to correlate with an increased yield based on initial decellularized ECM mass, where the iECM yield from aorta was significantly higher than all other batches (Fig 8e). This DF-TFF new method will therefore enable further evaluation of potential tissue specific differences and how they affect iECM bioactivity.

## Discussion

We previously developed iECM as a new type of ECM biomaterial to treat conditions characterized by excessive inflammation, including MI^7,10,11^, MODS^9^, and TBI^8^. We have demonstrated that this material is able to exert immunomodulatory and pro-reparative effects as seen by reductions in vascular permeability^7–9,11^, regulation of the immune system as seen by cytokine and gene expression^7,9,11^, improvements in cardiac function in small and large animal models of MI^7^, and improvements in survival in a mouse model of MODS^9^. While this material has shown great potential, the C-D-F method used to produce iECM had several flaws inherent to the manufacturing steps and as such limited its clinical translation, including limited manufacturing output and iECM concentration, as well as poor compliance with cGMP guidelines. For one, centrifugation limits scale up of iECM manufacturing due to volume processing limitations inherent to a given centrifuge model, necessitating expensive equipment upgrades for scaled up production. Even if more volume can be processed, the concentration of iECM through desalting and lyophilization requires a significant amount of time, further limiting production output. With regard to iECM concentration, centrifugation is unable to fully separate out all the >220 nm components, with an additional separation filtration step required. Because of this, the final concentration of iECM was limited since higher concentrations were difficult to pass through the filter. Finally, the use of dialysis tubing for concentrating isolated ECM components requires open-air human handling of the solution, increasing the potential for contamination and requiring handling loading of dialysis tubing in a highly clean environment, such as a laminar flow hood in order to be compliant with cGMP^12,13^. Altogether, the previously developed method for isolating the infusible fraction of pepsin digested ECM was effective at small scale manufacturing but contained many issues that limited efficient manufacturing scale up and clinical translation.

Our new DF-TFF addresses all of these limitations, paving the way for future iECM regulatory filings and clinical translation. When comparing iECM produced from 1 g digestions of decellularized ECM, the DF-TFF method produced iECM in a fraction of the time required by the C-D-F method. Furthermore, iECM produced with both methods were comparable, showing equivalent protein/peptide size distributions, biochemical properties, cytocompatibility, hemocompatibility, and binding to dysfunctional vasculature in the infarcted myocardium in a rat MI model. These comparisons suggest that the reductions in production time did not come with compromises in material properties, and iECM produced with DF-TFF was equivalent to iECM produced with C-D-F. Through the reduction in iECM production time afforded by the DF-TFF method, we were also able to process a digestion of 4g of decellularized ECM in a timely manner without significant change to the size distribution of iECM protein/peptides. The DF-TFF process thereby allowed for significant scale-up, which is important for clinical translation.

Due to the increased separation efficiency of DF and the concentration of iECM through TFF, we were also able to produce iECM of higher concentration than previously possible. Due to the inefficient separation of undesired large components by centrifugation, a secondary membrane filtration was required for full isolation of iECM components. This occurred after dialysis and lyophilization and required resuspension of the dry material at concentrations higher than the final concentration to account for losses during filtration. The upper bounds of this concentration were limited, as concentrations too high were difficult to filter. As such, through the use of the DF-TFF method, we are now able to produce iECM at higher concentrations. Leveraging these improvements in production, we were able to demonstrate iECM dose dependent efficacy in a porcine acute MI model and a mouse MODS model. While previous studies were limited to 4 mg/mL doses^7,9,11^, we were able to increase our dose to 6 and 8 mg/mL through improvements in iECM production. However, while 8 mg/mL iECM was possible to fabricate, the starting material required is substantial and concentrating to physiological salt levels is difficult due to hold-up volumes inherent to the TFF process. We thus chose to test this concentration in only 2 animals in the porcine MI study given the difficulties of producing 8 mg/mL at scale. Regardless, 6 mg/mL iECM still resulted in significant improvements in apical infarct angle, GWMI, and EDV with trending improvements in ESV, demonstrating the potential for improvements in therapeutic function enabled by DF-TFF. We only evaluated measures of infarct size, wall motion, and cardiac function at 24 hours to enable the same pigs to be used for retention as iECM is largely degraded by 3 days. While we did not perform echocardiography 24 hours post treatment in our previous porcine MI study, our previous rat MI study showed significant reductions in LV volumes at 24 hours that were maintained at 5 weeks post treatment^7^, suggesting these early changes we see with increasing dosage in this new porcine study could be maintained long term. Collectively, this therefore justifies performing a dose escalation study evaluating both 4 and 6 mg/mL in a first-in-man trial in acute MI patients.

Previously, in the rat MI model we found increased retention of iECM from 2.4 to 4 mg/mL^7^; however, interestingly, there was no increase in infarct retention of iECM between 4 and 6 mg/mL despite functional improvements, suggesting that other mechanisms beyond a local effect may be driving the increased efficacy. Crosstalk between the systemic immune system and heart has been discovered to play a significant role in disease progression in recent years^35–37^ and may also be playing a role with iECM. The increasing bioactivity we saw with increasing dose of iECM in the porcine MI model was also corroborated through our testing in the mouse MODS model, where the degree to which the expression of *Il6* was downregulated in the lungs also followed a linear dose response where increasing concentrations of iECM further reduced *Il6* expression. However, the local response with changes in vascular permeability in the MODS model was not impacted by increasing the dose from 4 to 6 mg/mL, again suggesting iECM may impact the systemic response in addition to having a local response. While these studies suggest increasing iECM concentration will be more efficacious from a translational standpoint, they also warrant new preclinical studies evaluating the systemic impact of iECM.

Finally, one of the major obstacles to further understanding and improving iECM as a biomaterial was the length of time required for batch production. While technically feasible to study the effect of tissue source on iECM’s material properties using the C-D-F method, the time required to produce multiple batches of iECM from multiple tissues is prohibitively long. Again, due to the improvements in manufacturing time afforded by the DF-TFF process, we were able to demonstrate the production of multiple batches of iECM from decellularized cardiac, lung, skeletal muscle, and aortic tissue in a timely manner. This preliminary work shows potential differences in protein/peptide size distribution and total material yield from iECM, again laying the foundation for further work evaluating how tissue specificity and composition may impact iECM bioactivity.

## Conclusion

In this study, we demonstrate the use of a novel method for processing decellularized ECM and manufacturing iECM involving DF-TFF. This method produces iECM comparable to previously reported batches with regard to material properties and *in vivo* targeting capacity while also significantly reducing production time and open-air human handling, improving compliance with cGMP guidelines. DF-TFF also enables increased production scale and increased iECM concentration, enabling preclinical small and large animal testing that demonstrates dose dependent bioactivity beyond what was previously achievable. Reduced production time also now enables timely production of iECM batches from alternate tissue sources, opening up the possibility of testing how changes in iECM composition can impact bioactivity. Overall, our DF-TFF iECM manufacturing method significantly reduces production time and open-air manipulation of liquids while also enabling increased iECM concentration, allowing for increased manufacturing scale, increased treatment efficacy and compliance with cGMP guidelines for further preclinical studies and future clinical translation.

## Methods

### 5.1: Decellularization of porcine tissue

Decellularization of porcine tissue was performed according to previously published protocols. Porcine hearts, lungs, aorta, and skeletal muscle (Collagen Solutions) were frozen immediately following harvest. Prior to decellularization, porcine heart, lungs, and skeletal muscle were minced using a meat grinder (LEM Products) and passed through a 3/16 inch metal grate for heart and lung tissue and a 1/4 inch metal grate for skeletal muscle. Porcine aorta was removed of fat and sliced into 1 cm squares. Cardiac, lung, and aorta tissue were decellularized using a solution of 1% SDS and phosphate buffered saline, (PBS), switching the solution 4 times over the course of 3 days. Skeletal muscle tissue was decellularized using a solution of 1% SDS and PBS, switching the solution 1 time over the course of 2 days. After decellularization, all tissues were rinsed with deionized water, rinsing for 2 hours and switching out the rinse solution 4 times. Skeletal muscle underwent an additional wash in isopropyl alcohol to remove fat tissue. Following decellularization, tissue was rinsed with water to remove residual detergent, lyophilized, milled, and passed through a 40-mesh metal grate.

### 5.2: Digestion of decellularized ECM

1000 mg or 4000 mg of milled ECM was suspended at 10 mg/mL of 1 mg/mL pepsin (≥2,500 units/mg protein, Millipore Sigma) dissolved in 0.1 M HCl. The ECM suspension was stirred with a stir bar for 48 hours at room temperature. After digestion, the ECM solution was processed into iECM using the below methods.

### 5.3: Preparation of iECM using centrifugation, dialysis, and filtration (C-D-F) method

To prepare iECM using the C-D-F method, digested ECM solution was neutralized and salt balanced according to previously published methods^7–9,11,17^. Digested ECM solution was first neutralized through the addition of 1 M NaOH. The salt balance was then corrected to 1X PBS through the addition of 20X PBS. Finally, the solution was diluted to 6 mg/mL ECM through the addition of 1X PBS. To separate out large ECM particulate matter, digested ECM was loaded into 50 mL conical tubes and centrifuged for 45 minutes at 15000 RCF. The supernatant was then removed and replaced with 1X PBS. The conical tubes were then vortexed to resuspend the pellet in the fresh PBS and the tubes were subjected to centrifugation at the same speed and time. To correct the protein and salt concentration to 4 mg/mL iECM and 1X PBS, the new supernatant was then combined with the previous supernatant and were loaded into cellulose ester dialysis tubing with a molecular weight cutoff of 1 kDa (Repligen). Supernatants were dialyzed against ultrapure water for 48 hours with 4 exchanges of the dialysis buffer occurring every 8-16 hours. After dialysis, ECM supernatant was transferred from dialysis bags to beakers, frozen, and lyophilized. After bulk lyophilization was completed, ECM supernatant was resuspended at approximately 16 mg/mL in 1X PBS. The proteins were further separated by size by filtration through a 0.45 μm filter and then sterile filtered through a 0.22 μm filter. The resulting solution now contained all components of iECM which was then aliquoted, lyophilized, and kept frozen until use. The mass of iECM in each aliquot was calculated by determining the difference in dry mass of the aliquots compared to the empty vial, then subtracting out the salts due to PBS.

### 5.4: Preparation of iECM using depth filtration and tangential flow filtration (DF-TFF) method

To prepare iECM through the DF-TFF method, digested ECM was first diluted to 2 mg/mL with a salt concentration of 0.5X PBS through the addition of 20X PBS and ultrapure water. Celpure P1000 (MilliporeSigma) was added to the digestion flask at a feed ratio of 60 g/L and the resulting suspension was stirred at 300 RPM for 30 minutes. After stirring, the suspension was filtered through a 20 μm paper filter (Whatman) under vacuum. The resulting Celpure filter cake was then rinsed with 200 mL 0.5X PBS. Both filtrates were combined and filtered through a 0.22 μm filter, resulting in dilute iECM components. iECM was then concentrated through recirculation tangential flow filtration against an open channel regenerated cellulose ultrafiltration cassette with a molecular weight cutoff of 1 kDa (Pellicon). iECM was concentrated to 1/10 the original volume, after which it was terminally sterile filtered through a 0.22 μm filter, aliquoted, lyophilized, and kept frozen until use. The mass of iECM in each aliquot was calculated by determining the difference in dry mass of the aliquots compared to the empty vial, then subtracting out the salts due to PBS.

### 5.5: Characterization of iECM protein/peptide size distribution

iECM protein/peptide size distribution was quantified by first resuspending at 4 mg/mL. 200 μL of iECM was separated through an ENrich SEC 650 size exclusion column (Bio-Rad) with a separation range of 5-650 kDa and detected through absorbance at 215 nm^11^. The molecular weights of iECM proteins were estimated by comparing the peaks to a standard curve generated from a protein gel filtration standard (Bio-Rad). Relative ratio of the iECM protein/peptide sizes based on SEC peak was determined by integrating the peaks of the SEC trace and dividing the area of the individual peaks to the total area under the curve.

### 5.6: Biochemical characterization of iECM

#### 5.6.1: Quantification of residual double stranded DNA (dsDNA)

Residual dsDNA from the decellularization process was quantified through a fluorescent PicoGreen assay (Thermofisher). Residual dsDNA was first isolated from iECM using a Nucleospin DNA isolation kit (Machnery-Nagel) according to manufacturer protocols. Following isolation, the concentration of dsDNA was quantified using PicoGreen and Lambda DNA standards (Thermofisher). DNA content was then normalized to the mass of iECM loaded into each isolation column.

#### 5.6.2: Quantification of residual SDS

Residual SDS from the decellularization process was quantified through complexation with methylene blue. iECM samples were mixed with aqueous methylene blue solution (50 μg/mL in 1 mM Tris buffer) and vortexed to form methylene blue-SDS complexes. Methylene blue-SDS complexes were extracted by mixing with chloroform, with the mixtures being allowed to settle into two phases to extract the methylene blue-SDS complex into the organic layer. Concentration of SDS was determined by measuring the absorbance of the solution at 655 nm, with the concentration determined by comparing to a calibration curve of known methylene blue-SDS complexes. Residual SDS was normalized to the amount of iECM loaded into the assay.

#### 5.6.3: Estimation of iECM protein content

Protein content of iECM batches were estimated using solution absorbance of 215 nm light^38^., iECM batches were diluted by 10X and their absorbance was determined using a NanoDrop 2000 spectrophotometer (Thermofisher), with estimated protein content determined by comparing to a BSA standard.

#### 5.6.4: Quantification of sulfated glycosaminoglycan (sGAG) content

sGAG content of the iECM batches was determined through complexation with dimethylmethylene blue (DMMB)^7,11^. iECM was mixed with DMMB solution (32 μg/mL DMMB in formate buffer) at a ratio of 1:10 v/v. The solutions were mixed for 30 minutes to form DMMB-sGAG complexes, after which the vials were centrifuged for 10 minutes at 14000 RCF. DMMB-sGAG pellets formed after centrifugation and the supernatant was carefully aspirated to remove unbound dye. DMMB-sGAG complexes were decomplexed using 8M guanidine hydrochloride for 30 minutes. The concentration of DMMB that was previously bound to sGAGs was determined by measuring the absorbance at 656 nm, allowing for quantification of sGAGs in iECM. Concentrations of sGAGs were determined by comparing to a calibration curve of known concentrations of chondroitin sulfate (Thermofisher).

#### 5.6.5: Quantification of free amine content

iECM free amine content was determined through a fluorometric o-Pthaldialdehyde reagent assay. O-Pthaldialdehyde reagent (Thermofisher) was mixed directly with iECM diluted by 100X at a ratio of 10:1 in 96 well plates. After 5 minutes of mixing, fluorescent intensity of wells was determined using a M200 Infinite plate spectrophotometer (Tecan). Free amines of iECM were determined by comparing to a glycine standard.

### 5.7: Characterization of iECM Viscosity

To determine the viscosity of iECM as a product of manufacturing process, C-D-F iECM at 4 mg/mL and DF-TFF iECM at 4 mg/mL were tested using a TA Instruments Discovery Hybrid Rheometer equipped with a 20 mm parallel plate geometry. Viscosity was determined by measuring shear stress across a flow sweep from 1 to 1000 Hz at 25°C.

### 5.8: Cytocompatibility of iECM

Cytocompatibility of iECM batches was determined by measuring their effect on the metabolic activity of rat coronary artery endothelial cells (rCAECs). rCAECs were cultured to 70-80% confluency in endothelial cell growth medium (Promocell) supplemented with 1% penicillin/streptomycin. Cells were then detached using 0.25% trypsin/EDTA (Gibco), counted, and seeded in 96 well plates at 10,000 cell/well. Cells were allowed to attach for 24 hours, after which the supernatant was removed and replaced AlamarBlue (Thermofisher) diluted in each cell’s respective media. Cells were incubated with AlamarBlue for 1 hour, after which the supernatant was removed and analyzed using a spectrophotometer to determine the baseline metabolic rate. Supernatants were then replaced with 0.4 mg/mL C-D-F iECM, 0.4 mg/mL DF-TFF iECM, or PBS. Treatments were diluted using the appropriate cell media for the cell types being used. As a negative control, cells were treated with 70% ethanol. Cells were incubated with treatments for another 24 hours, after which supernatants were removed and replaced with AlamarBlue. Cells were incubated again for 1 hour and supernatants were analyzed to determine post-treatment metabolic rate. Cytocompatibility was determined by determining changes in metabolic rate and comparing to positive and negative controls.

### 5.9: Hemocompatibility of iECM

Hemocompatibility of iECM batches with human blood samples (n = 4) was determined by assessing their effects on red blood cell (RBC) and platelet aggregation according to previously published protocols^7,23,24^. iECM was diluted 1:10 prior with human blood to testing, with this dilution representing the physiologically relevant dilution that would occur following intracoronary infusion. Red blood cell aggregation was measured with a photometric rheoscope (Myrenne Aggregometer, Myrenne) after adjusting the hematocrit to 45% with autologous plasma. Platelet aggregation was determined using isolated platelet rich plasma with a lumi-aggregonometer (Chrono-log), with platelet coagulation cascade agonists adenosine diphosphate (ADP), epinephrine, and collagen added (1:200 – 1:1000 dilutions).

### 5.10: Assessment of iECM in rodent models

All procedures in this study were approved by the Institutional Animal Care and Use Committee at the University of California San Diego and in accordance with the guidelines of the Association for the Assessment and Accreditation of Laboratory Animal Care (A3033-01).

#### 5.10.1: Retention of iECM in a rat MI model

To induce MI, adult female Sprague-Dawley rats (225-250g) were used as previously described^7,10,11^. To induce acute MI, the left coronary artery was accessed via a thoracotomy and ligated for 35 minutes with a suture. Following ischemia, the suture was released to restore blood flow. For simulated intracoronary infusion, the aorta was temporarily clamped and the 200 μL of treatment was injected into the LV lumen using a 30 G needle. Infusions were performed within 10 minutes of reperfusion. Animals were randomly assigned treatment groups.

To determine biodistribution and LV localization of iECM based on manufacturing method, iECM was labeled with VT-750 at a ratio of 1:100 by volume. iECM was labeled prior to fractionation with ethanol to preserve the relative labeling ratio found in complete iECM. Labeled material was delivered in the rat model of MI as previously described^11^. At time points of 1 hour, rats were euthanized via an overdose of sodium pentobarbital via intraperitoneal injection and perfused with PBS to remove residual iECM in the blood. Heart and satellite organs (lungs, kidneys, spleen, liver) were extracted, with the heart sliced into 6 transverse slices. Heart slices and satellite organs were scanned with an Odyssey CLx (LICORbio) near infrared scanner to determine total signal.

#### 5.10.2: Effect of iECM on vascular permeability in a mouse MODS model

iECM effects on vascular permeability in a mouse MODS model was tested as previously published^9^. 9-week-old wild type female C57BL/6J mice (Jackson Laboratories) received intraperitoneal (IP) injections of 1 mg/mL lipopolysaccharide in sterile saline (LPS) for an initial dose of 10 mg/kg. After 6 hours, mice received an additional 1 mg/kg dose of LPS. After 10 hours post initial LPS injection, mice received an intravascular injection of 200 µL sterile saline, C-D-F iECM, or DF-TFF iECM through the tail vein. 1 hour post saline or iECM delivery, mice received an intravenous injection of 200 μL 10 mg/mL BSA labeled with 10 mg/mL IVISense 750 NHS fluorescent dye (Revvity) at a ratio of 1:100 v/v. After 1 hour, mice were euthanized, perfused with 1X PBS, and lungs, heart, spleen, kidneys, and brains were harvested. Near infrared signal of organs was scanned using an Odyssey CLx near infrared scanner (LICORBio). Average fluorescent intensity was normalized to the average signal of saline treated animals.

For assessing *Il6* gene expression, 9-week-old wild type female C57BL/6J mice under went the same LPS injection and iECM administration as above. At 30 hours post initial injection, mice were euthanized and lungs were harvested. mRNA was isolated from lung tissue and converted to cDNA. Quantitative PCR (qPCR) was used to determine relative fold expression of *Il6* in iECM treated animals compared to *Gapdh* and saline using the ΔΔCt calculation method.

### 5.11: Testing of iECM in a porcine acute MI model

All procedures in this study were approved by the Institutional Animal Care and Use Committee at CRF Skirball Center for Innovation. Procedures were conducted at a test facility accredited by the Association for the Assessment and Accreditation of Laboratory Animal Care and registered with the United States Department of Agriculture.

To test retention and effects on cardiac function of DF-TFF iECM in a large animal model, female Yucatan mini swine (Premier Biosource) between 9.6 and 14.6 months of age underwent MI through occlusion of the left anterior descending (LAD) artery. Vascular access was first obtained through the placement of a vascular access sheath in the left jugular vein and the left carotid artery. Myocardial ischemia was induced through balloon occlusion of the mid LAD for 90 minutes, with the balloon deflated to allow for reperfusion. Following reperfusion, 4 mL iECM or saline was infused at 1 mL/min with a 2 min-stop flow, with 2 infusion cycles being performed for full treatment delivery. Prior to infusion, iECM was fluorescently labeled with Ivisense-750 as described above for detection using near infrared scanning. After 24 hours, blood was collected for hematology, serum chemistry, and coagulation analysis. Animals were then euthanized and 6 x 4 mm sections were taken from the infarct region. Sections 3 and 5 were scanned on an Odyssey 9120 (LICORbio) to assess retention of iECM in the infarcted myocardium.

Transthoracic 2D echocardiography was performed using a Philips EPIQ 7C Ultrasound system with X5-1 Transducer. Measurements were performed prior to ischemia as a baseline, as well after reperfusion to ensure an adequate MI, after intracoronary infusion, and prior to euthanasia at 24 hours. Echocardiography was used to estimate infarct area using the angle of detected left ventricle wall motion abnormality. End diastolic volume (EDV), end systolic volume (ESV), and ejection fraction (EF) were derived from echocardiography images, with changes in cardiac function determined by assessing percent change between terminal values and post-MI (pre-treatment) values. Myocardial infarction was confirmed by an increase in intraventricular septum thickness, an observed infarction angle, and an elevated GWMI. A total of 18 animals underwent the MI procedure, with 3 animals dying during occlusion and 2 animals excluded from further analysis due to lack of evidence for MI (no detected infarct angle and no increase in GWMI). To prevent measurement bias, personnel were blinded to treatment groups for measurement and data analysis.

### 5.12: Statistical Analysis

Statistical analysis was performed using GraphPad Prism. For analysis of 2 groups, statistical significance was determined using a 2-tailed unpaired t-test. For analysis of 3 or more groups, a one-way ANOVA with post hoc Tukey or Dunnett test was performed to determine statistical significance. Significant dose response of iECM was determined using a linear trend analysis. Significance was accepted at p<0.05.

## Acknowledgements

This work was supported in part by the California Instititute for Regenerative Medicine (TRAN1-15291) and the NIH NHLBI (R01HL165232 to KLC and R44HL169072 to AK). MN was funded by an NHLBI training grant (T32HL0074444) and an AHA Postdoctoral Fellowship (24POST1242447). We would also like to thank our manufacturing partners for their input for the development of the DF-TFF process. We would also like to thank Dr. Brent Stockwell and Mars Fongheiser at Columbia for kindly allowing us to use their Licor Odyssey 9120. Timeline graphics in figures were made using Biorender.com KLC is co-founder, consultant, board member of, and holds equity interest and receives income from Ventrix Bio Inc. EZ is a consultant for and receives income from Ventrix Bio Inc.

