## Supplemental Information for "An Improved Manufacturing Method for Infusible Extracellular Matrix Materials to Enable Efficient Scale-Up, Improved Efficacy, and Clinical Translation"

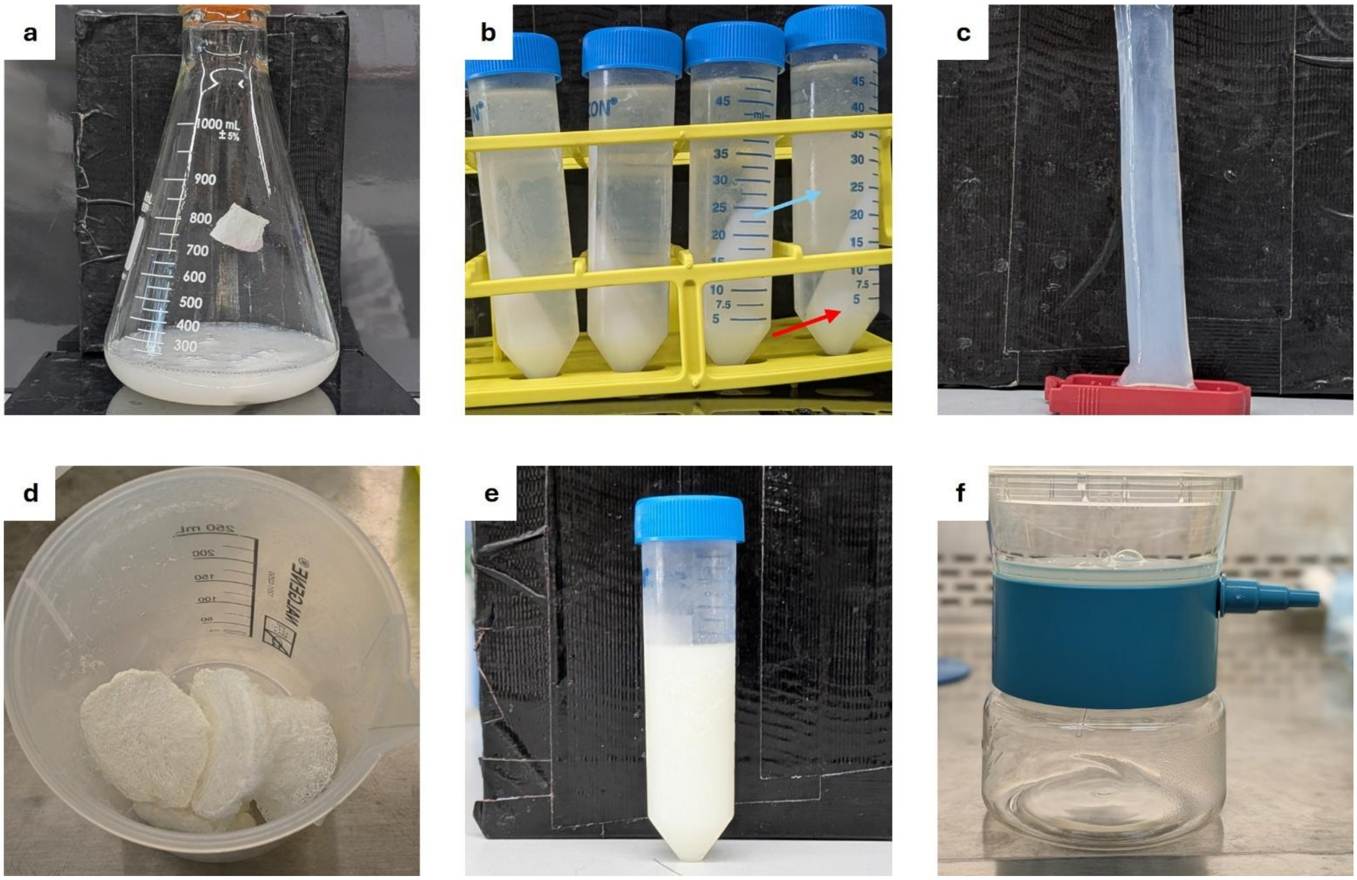


**Figure S1: Photos of C-D-F iECM manufacturing process.** For fabricating iECM using the C-D-F manufacturing method, decellularized ECM powder is first digested with pepsin (a) then centrifuged (b) to separate large particulate matter (red arrow) from lower molecular weight components (blue arrow). Supernatant is then dialyzed (c) to desalt prior to lyophilization (d) and resuspension at a higher concentration (e). Prior to terminal sterile filtration, residual large ECM particles are separated through a membrane filtration (f).

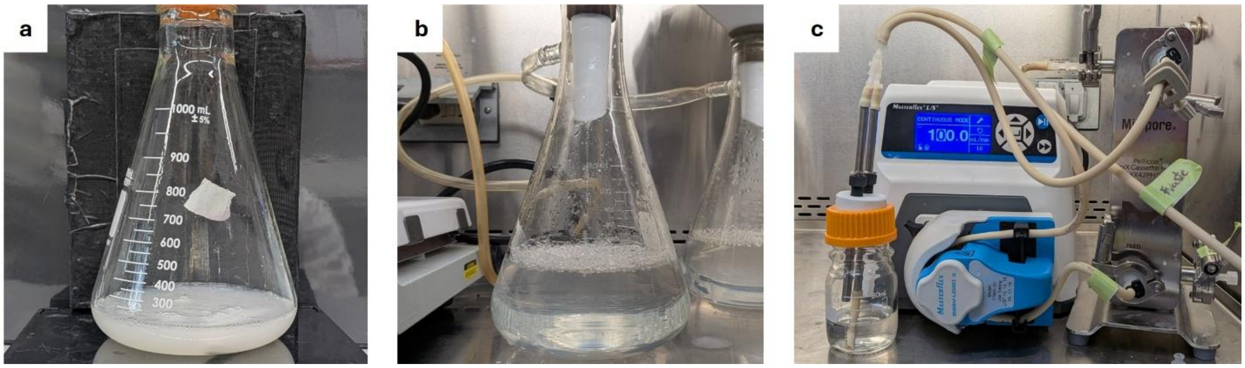


**Figure S2: Photos of DF-TFF iECM manufacturing process.** To produces iECM using the DF-TFF method, decellularized ECM is first digested with pepsin (a), then diluted and combined with diatomaceous earth filter aid prior to filtration (b). Filtered iECM was then concentrated using tangential flow filtration (c) prior to terminal sterile filtration.

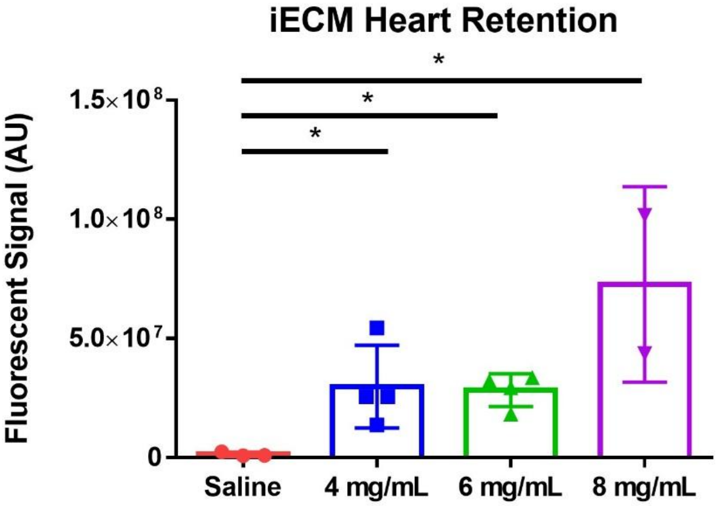


**Figure S3: Retention of iECM in pig hearts with increasing iECM dose as determined by total fluorescent signal.** Statistical significance ($p<0.05$) determined with a one-way ANOVA with post-hoc Dunnett test compared to saline (*$p<0.05$)

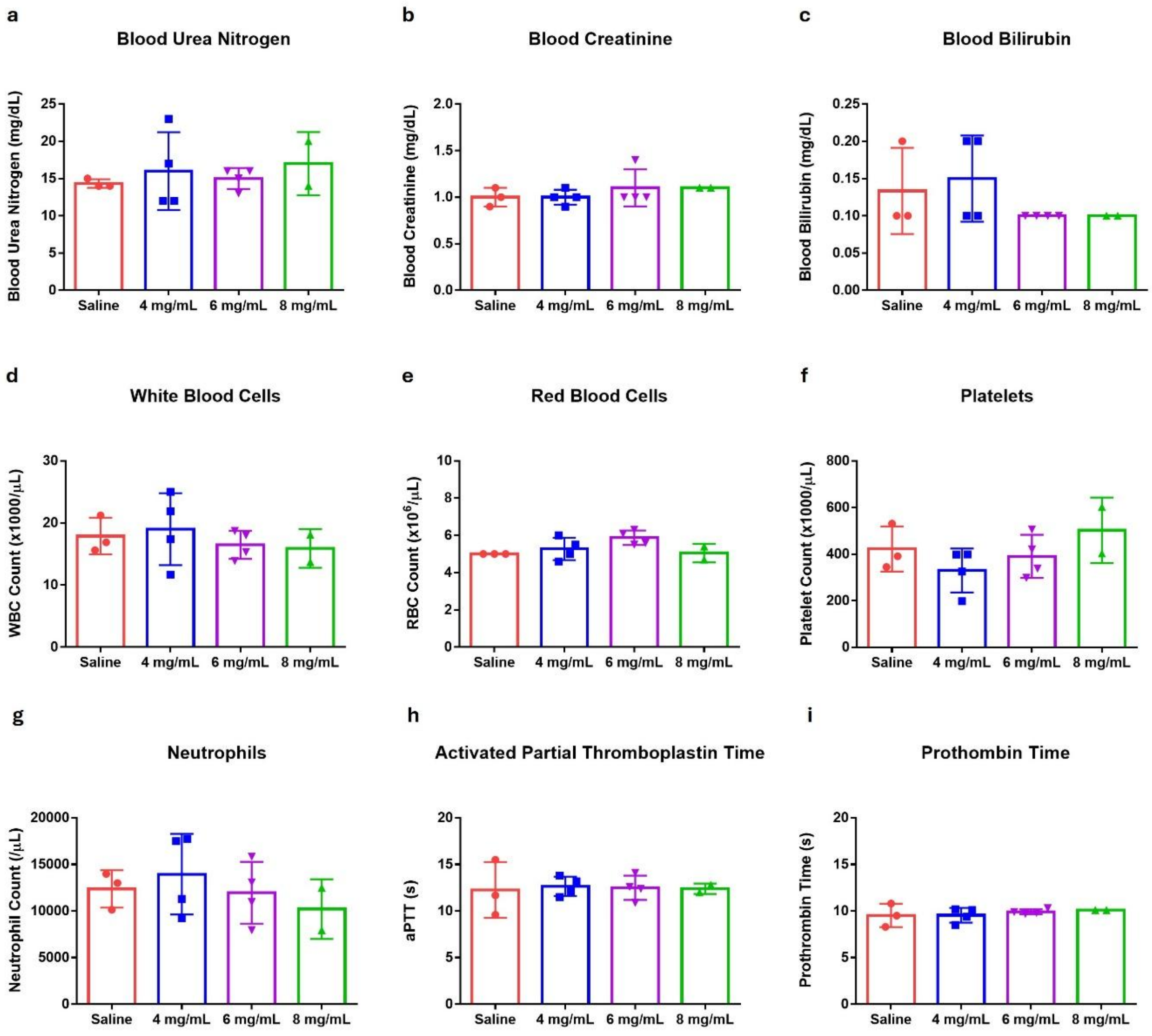


**Figure S4: Clinical pathology data showing safety of increased iECM doses compared to saline.** a) Blood urea nitrogen levels of pigs 24 hours post treatment infusion. b) Blood creatinine levels of pigs 24 hours post treatment infusion. c) Blood bilirubin levels of pigs 24 hours post treatment infusion. d) Total white blood cell count of pigs 24 hours post reperfusion. e) Red blood cell count of pigs 24 hours post reperfusion. f) Platelet count of pigs 24 hours post reperfusion. g) Neutrophil count of pigs 24 hours post reperfusion. h) Activated partial thromboplastin time of blood collected from pigs 24 hours post reperfusion. i) Prothrombin time of blood collected blood collected from pigs 24 hours post reperfusion. All tests showed no significant difference compared to saline, demonstrating no adverse effects associated with increased iECM concentration. Statistical significance ($p < 0.05$) determined with a one-way ANOVA.
